# Biochemical Pathways Represented by Gene Ontology Causal Activity Models Identify Distinct Phenotypes Resulting from Mutations in Pathways

**DOI:** 10.1101/2023.05.22.541760

**Authors:** David P Hill, Harold J Drabkin, Cynthia L Smith, Kimberly M Van Auken, Peter D’Eustachio

## Abstract

Gene inactivation can affect the process(es) in which that gene acts and causally downstream ones, yielding diverse mutant phenotypes. Identifying the genetic pathways resulting in a given phenotype helps us understand how individual genes interact in a functional network. Computable representations of biological pathways include detailed process descriptions in the Reactome Knowledgebase, and causal activity flows between molecular functions in Gene Ontology-Causal Activity Models (GO-CAMs). A computational process has been developed to convert Reactome pathways to GO-CAMs. Laboratory mice are widely used models of normal and pathological human processes. We have converted human Reactome GO-CAMs to orthologous mouse GO-CAMs, as a resource to transfer pathway knowledge between humans and model organisms. These mouse GO-CAMs allowed us to define sets of genes that function in a causally connected way. To demonstrate that individual variant genes from connected pathways result in similar but distinguishable phenotypes, we used the genes in our pathway models to cross-query mouse phenotype annotations in the Mouse Genome Database (MGD). Using GO-CAM representations of two related but distinct pathways, gluconeogenesis and glycolysis, we show that individual causal paths in gene networks give rise to discrete phenotypic outcomes resulting from perturbations of glycolytic and gluconeogenic genes. The accurate and detailed descriptions of gene interactions recovered in this analysis of well-studied processes suggest that this strategy can be applied to less well-understood processes in less well-studied model systems to predict phenotypic outcomes of novel gene variants and to identify potential gene targets in altered processes.

**Summary:** Genes act in interconnected biological pathways, so single mutations can yield diverse phenotypes. To use the large body of mouse functional gene annotations, we converted human Gene Ontology-Causal Activity Models (GO-CAMs) of glucose metabolism to orthologous mouse GO-CAMs. We then queried phenotypes for mouse genes in these GO-CAMs and identified gene networks associated with discrete phenotypic outcomes due to perturbations of glycolysis and gluconeogenesis. This strategy can be extended to less well-understood processes and model systems to predict phenotypic outcomes.

## Introduction

Gene products mediate networks of causally connected functions to give rise to biological outcomes, revealed as observable phenotypes. Phenotypes correlate with gene functions, and GO Biological Processes can be predicted by analyzing phenotypic outcomes (Ascensao *et al*. 2014). Causally connected gene functions can be grouped into biological pathways that represent their co-ordinated action to achieve a specific outcome. However, phenotypes, even ones conventionally attributed to a single mutated gene, can be the result of interactions of multiple genes. Phenotypic variability among affected individuals described as incomplete penetrance and variable expressivity can be the result of interactions between the mutated gene and other genes with which it normally interacts (Kingdom and Wright 2022).

Genes can also display epistatic interactions in which the phenotype caused by a mutation in one gene can mask a phenotype of another variant gene in the same pathway (de Visser *et al*. 2011). Recent studies in large human populations of the range of phenotypes observed in individuals heterozygous for CFTR (cystic fibrosis transmembrane conductance regulator) mutations (Barton *et al*. 2022), mutations in genes associated with developmental delay (Kingdom *et al*. 2022), and mutations in genes associated with maturity onset diabetes of the young (Mirshahi *et al*. 2022) provide striking illustrations of the complex interactions that occur between a mutation in one gene and normal genomic polymorphisms resulting in an affected individual’s phenotype.

Pathway databases, which model biological processes as networks of reactions, provide useful tools for exploring such interactions. An individual reaction converts input physical entities into modified or relocated output entities. The conversion is mediated by regulatory and catalytic activities of still other entities. A reaction is thereby temporally and causally connected to other reactions that generate its inputs, consume its outputs, or modulate the activities of its catalysts and regulators, and loss of a single protein’s function can result in multiple phenotypic consequences determined by the reaction network structure. Understanding this structure thus provides a framework for interpreting phenotypes associated with mutations in genes present in the network.

How generally can this network model be applied to predict phenotypes resulting from the disruption of individual genes given diverse genetic backgrounds? Can we use phenotypes to tease apart subpathways in a network and identify the ones whose disruption gives rise to a discrete phenotype? Here, we used a network of pathways representing glucose and pyruvate metabolism to address these questions. Under normal physiological conditions in vertebrates, glucose levels are tightly controlled despite variation in demands for energy and biosynthetic intermediates that depend on glucose metabolism, and variation in glucose availability from dietary intake, glycogen breakdown and de novo glucose synthesis.

The complementary, interconnected processes of glycolysis and gluconeogenesis, in which glucose is catabolized and synthesized, respectively, are central to glucose homeostasis. Glycolysis converts glucose and other monosaccharides to pyruvate and generates ATP and reduced NAD (Prochownik and Wang 2021). Gluconeogenesis consumes pyruvate, NADH, and ATP to generate glucose (Chourpiliadis and Mohiuddin 2022). Lactate (Cori 1981) and amino acids (Felig 1973) are major sources of carbon skeletons that are converted to pyruvate, and there are three routes from pyruvate to a common phosphoenolpyruvate intermediate (Figure 1A). From phosphoenolpyruvate to glucose is a ten-reaction process. Seven reactions are shared by gluconeogenesis and glycolysis: these reactions are reversible under physiological conditions, mediated by the same enzymes, and directed by mass action. The remaining reactions of each process are irreversible under physiological conditions and mediated by different enzymes.

**Figure 1.**
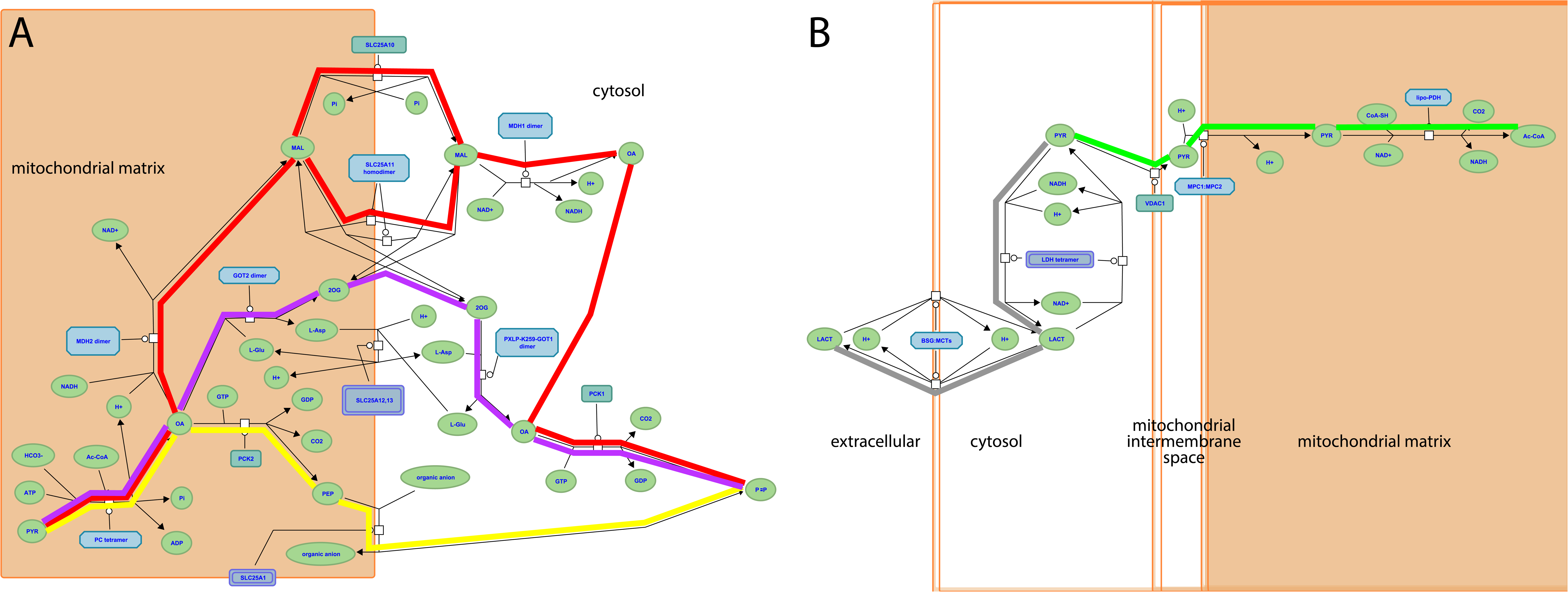
**Reactome pathways. A**, three alternative routes from mitochondrial pyruvate to cytosolic oxaloacetate at the start of the gluconeogenesis pathway (https://reactome.org/PathwayBrowser/#/R-HSA-70263), corresponding to GO-CAM models Gluconeogenesis 1 (red highlighting), 2 (yellow), and 3 (magenta). **B**, aerobic conversion of pyruvate to acetyl-CoA (green highlighting) and anaerobic conversion to lactate (gray) (https://reactome.org/PathwayBrowser/#/R-HSA-70268). In each case, the process description format shows the substrates and products (chemical entities) of each reaction, the identities of the proteins or complexes that mediate it, and causal connections due to shared products / substrates.

Glycolysis occurs in all tissues in the body. For red blood cells, which lack mitochondria, glycolysis is the only source of ATP. In contrast, gluconeogenesis is mainly confined to hepatocytes and kidney proximal tubules. The irreversible steps of glycolysis and gluconeogenesis are normally coordinately regulated to maximize hepatocyte glycolysis when blood glucose levels are high and in skeletal muscle in response to strenuous exercise, and to minimize hepatocyte glycolysis and maximize hepatocyte and kidney proximal tubule gluconeogenesis in response to fasting and physiological stresses. The pyruvate end-product of glycolysis is metabolized aerobically to acetyl-CoA or anaerobically to lactate (Figure 1B).

We distinguished three groups of genes and regulatory processes for our analysis: those unique to glycolysis, those unique to gluconeogenesis, and those shared between glycolysis and gluconeogenesis, allowing us to ask whether variant forms of genes in each set are associated with distinct or shared phenotypes. Shared phenotypes for genes in one set that are distinct from phenotypes of another set, would allow us to associate an individual biochemical pathway with specific outcomes. Thus we could identify genes in the pathway that could potentially be manipulated to modify clinical outcomes.

We have previously computationally generated GO-CAM models of the human processes of glycolysis and gluconeogenesis, and of the metabolism of pyruvate, the end-product of glycolysis, to lactate (anaerobic metabolism) or acetyl-CoA (aerobic metabolism) (Good *et al*. 2021). The GO paradigm annotates individual gene products with terms that describe their molecular functions, association with cellular components, and involvement in multistep biological processes (The Gene Ontology Consortium 2017). GO-CAM models link these annotations of individual gene products into a causal flow. In GO-CAM models of metabolic processes, this causal flow is represented by enzymatic activities enabled by gene products that are connected by shared input, output molecules or regulatory functions of gene products within the causal chain (Thomas *et al*. 2019). A GO-CAM differs from the Reactome model of the same process in that the latter employs a more discursive process description data model (Le Novère *et al*. 2009) to describe processes as networks of reactions mediated by gene products that transform input molecules into output molecules (Gillespie *et al*. 2022). GO-CAM models derived from Reactome pathways can be used to define sets of genes that function in linear causal pathways and can be used to discriminate genes that function in multiple or branched pathways. A specific feature of GO-CAM models generated in the work described here is that each covers a group of molecular functions corresponding to a single GO biological process, while Reactome models are more variable in size and scope, combining, for example, metabolic and regulatory functions, and merging multiple tissue-specific variant forms of a process.

We have now used computationally-derived human GO-CAM models as templates to manually construct GO-CAM models of the corresponding mouse pathways, teasing apart pathway networks to create a causal flow of GO molecular functions enabled by mouse gene products. We used the gene lists from these models as a basis to interrogate MGD phenotype annotations to determine if mutant forms of mouse genes associated with different mouse GO-CAM models were enriched for different phenotypes (Blake *et al*. 2021). This approach takes advantage of the wealth of data in the laboratory mouse linking genetic variation to phenotypic variation (http://www.informatics.jax.org/greenbook/).

Laboratory mice are a uniquely rich resource of mammalian phenotypes associated with variations in single genes expressed on diverse genetic backgrounds. A large and diverse collection of mouse genetic variants associated with visible phenotypes has been assembled in the past 120 years by systematically searching for visibly abnormal individuals in inbred populations, by inducing random mutations by mutagens, radiation or insertion of gene trap constructs and by modern genome editing technologies such as targeted mutagenesis in ES cells or CRISPR/Cas9 methodologies (reviewed in Bello *et al*. 2021). The variant heritable alleles generated in these ways involve DNA sequence changes including point mutations, insertions of heterologous sequence, chromosomal deletions and rearrangements affecting several genes, and their phenotypes have been assessed to various extents, by individual investigators and in small and large scale mutagenesis projects (Groza *et al*. 2023). These disparate phenotype data whether generated by large projects, or by expert curation of published literature, are standardized using terms from the Mammalian Phenotype ontology (https://www.informatics.jax.org/vocab/mp_ontology) (Smith *et al*. 2005) and to relevant experimental strain genetic backgrounds and evidence, and are integrated in the phenotypes section of the Mouse Genome Database (MGD) resource

(https://www.informatics.jax.org/phenotypes.shtml) (Bello *et al*. 2015). MGD genotype-phenotype data include simple genotypes with causative mutations in a single gene as well as more complex genotypes where mutations in multiple genes may contribute to the phenotype. For analyses here only the simple genotype associations were used (Bello *et al*. 2016).

Lastly, we asked if we could extend our knowledge mechanistically by using enriched phenotypes for gluconeogenesis to identify transcription factors that are associated with the same phenotypes. We then confirmed that some of the transcription factors we identified are involved in regulation of gluconeogenesis while others are involved in pathways that interact with the gluconeogenic pathway. With this inferential approach, we can begin to systematically expand our pathway analysis to include coordinated transcriptional regulation of the pathway. Combining transcriptional regulation with well-studied allosteric regulation of the enzymes of the pathway, we can begin to better understand the causal connections among genes that result in complex abnormal phenotypes.

## Materials and Methods

### Genes Used in GO-CAM Models

GO-CAM (Thomas *et al*. 2019) activity flow (Le Novère *et al*. 2009) models computed from Reactome gluconeogenesis, glycolysis, pyruvate – lactate, and pyruvate – acetyl-CoA pathways (Gillespie *et al*. 2022; Good *et al*. 2021; Hill *et al*. 2016) assigned functions to 60 human gene products (Table 1). These gene products are grouped in Reactome into sets of paralogs that share a function (e.g., the “glucokinase” set consists of *GCK*, *HK1*, *HK2*, *HK3*, and *HKDC1*). We identified high-confidence mouse orthologs of these human gene products: mouse gene products with experimental evidence of conserved function from direct assays (Blake *et al*. 2021; Ringwald *et al*. 2022). Supporting evidence, such as references and evidence codes, was added to the mouse models using standard protocols for GO annotation (Balakrishnan *et al*. 2013; The Gene Ontology Consortium 2017). This search yielded 44 mouse gene products, typically one or two corresponding to each human set. This search failed for one, *Slc25a1* (MGI:1345283): despite its sequence similarity to human *SLC25A1* we found no experimental evidence to support its role in phosphoenolpyruvate transport in mice and only suggestive indirect evidence in related species. One mouse gene product with no human counterpart, *Eno1b* (MGI:3648653), was identified: this gene arose by retrotransposition after the divergence of mouse and human lineages from their last common ancestor (Bulusu *et al*. 2017). The 44 mouse gene products identified in this step were used to construct mouse GO-CAM models. To construct a mouse GO-CAM, each molecular function node in the corresponding human GO-CAM was populated with the one or two mouse gene products known from direct experimental evidence to enable that function.

The part of the Reactome gluconeogenesis model that represented alternative pathways from pyruvate to phosphoenolpyruvate was split into separate mouse GO-CAMs. Separating the Reactome pathways that represent multiple branches of a single process into individual GO-CAMS allows us to compare and contrast ‘subpathways’ as in our analysis of the fates of pyruvate described below. We created four pathways to represent glycolysis and gluconeogenesis: Canonical glycolysis 1 (Mouse) – gomodel:5745387b00001516, Gluconeogenesis 1 (Mouse) – gomodel:61e0e55600001225, Gluconeogenesis 2 (Mouse) – gomodel:62d0afa500000132, and Gluconeogenesis 3 (Mouse) – gomodel:62d0afa500000298 (Figure 1A and Table 1). For analyses of downstream pathways, we created two mouse GO-CAM models based on the Reactome ‘Pyruvate Metabolism’ (R-HSA-70268) pathway, one representing the interconversion of pyruvate and lactate – gomodel:633b013300001238) and one representing the conversion of pyruvate to acetyl-CoA – gomodel:633b013300001469) (Figure 1B). All GO-CAM models can be browsed at http://model.geneontology.org/##, where ## is replaced by the model ID (e.g., http://model.geneontology.org/633b013300001238).

### Genes Used for Phenotype Enrichment Analysis

We extended the list of mouse – human gene product pairs used to construct mouse GO-CAM models by searching MGD for all mouse orthologs of human set members supported by any experimental annotations or sequence orthology annotations consistent with conserved function. The resulting list contains 59 human – mouse pairs, excluding human *SLC25A1* (no experimentally verified mouse counterpart) and mouse *Eno1b* (no human counterpart) (Table 1).

The mouse gene products on this larger list were assembled into five sets that were derived from the GO-CAM models and represented genes whose functions are causally connected in each pathway. These GO-CAM-derived ‘causal’ gene sets were used for phenotype enrichment analyses: 1) genes unique to glycolysis, 2) genes unique to gluconeogenesis, 3) genes shared between glycolysis and gluconeogenesis, 4) genes unique to glycolysis plus genes that convert pyruvate to lactate, and 5) genes unique to glycolysis plus genes that convert pyruvate to acetyl-CoA. Enrichment analysis was performed on the gene sets using the VisuaL Annotation Display (VLAD) tool (http://proto.informatics.jax.org/prototypes/vlad/) (Richardson and Bult 2015) using default settings and MGD annotation data complete through January 26, 2023. For comparison of sets 1, 2, and 3 we used the comparative analysis functionality of VLAD. This functionality calculates an enrichment score for each set and plots the relative enrichment significance in graphical format overlaying the structure of the Mammalian Phenotype Ontology (MP) (Smith and Eppig 2012). Results are also generated in a tabular format. We hypothesized that genes causally connected in the separate branches would share separate phenotypes with the main branch. We analyzed sets 1, 4, and 5 individually. We filtered the results of each analysis using a false discovery rate (FDR) q<0.05 cut-off. If a phenotype term, e.g., MP:0013663 “increased myeloid cell number”, and a more specific is_a child of that term, e.g., MP:0002640 “reticulocytosis” both fell below the q<0.05 cut-off, we considered only the child term in our analyses here. Next, we determined which of these significant phenotypes had a greater number of genes associated with glycolysis and either downstream pathway compared with glycolysis alone or with the other downstream pathway. Complete results from all VLAD analyses are shown in supplemental tables s1-s5.

### Analysis of Transcription Factors That Correspond to Phenotypes Enriched in the Gluconeogenesis Gene Set

To identify transcription factors that are associated with phenotypes enriched in the gluconeogenesis genes, we searched the MGD ‘Genes and Markers Query Form’ on (https://www.informatics.jax.org/marker – February 2, 2023 data) for genes annotated to the GO term ‘DNA binding transcription factor activity’ (GO:0003700) and to the ten most specific enriched MP terms with a FDR q<0.05 in the VLAD graph that correspond to specific metabolic or developmental processes: ‘hypoglycemia’’ (MP:0000189), ‘increased liver triglyceride level’ (MP:0009355), ‘decreased circulating triglyceride level’ (MP:0002644), ‘increased liver glycogen level’ (MP:0010400), ‘increased kidney glycogen level’ (MP:0031003), ‘decreased granulocyte number’ (MP:0000334), ‘abnormal midbrain development’ (MP:0003864), ‘abnormal gluconeogenesis’ (MP:0003383), ‘increased circulating triglyceride level’ (MP:0001552), and ‘hepatic steatosis’ (MP:0002628).

### Data Access

Full results of our analyses are given in Tables 1-4 and in the supplementary files for this paper. The full Reactome and MGD data sets are freely available to all users at www.reactome.org and https://www.informatics.jax.org/, respectively.

## Results

### Creation and validation of mouse GO-CAM models

To determine whether we could associate biochemical pathways with specific phenotypes, we first built mouse GO-CAM models for glycolysis, gluconeogenesis, and pyruvate metabolism. Reactome-derived human GO-CAM models served as templates, and were populated with orthologous mouse gene products available from MGD (Table 1). We generated a single mouse pathway for glycolysis. For gluconeogenesis we generated three mouse GO-CAM models, one for each of the three distinct routes to the formation of cytosolic oxaloacetate in the Reactome model of human gluconeogenesis (https://reactome.org/PathwayBrowser/#/R-HSA-70263), (Figure 1A, Table 1). We generated models for anaerobic metabolism of pyruvate to lactate and aerobic metabolism to acetyl CoA, representing branches of pyruvate metabolism in the Reactome human model (https://reactome.org/PathwayBrowser/#/R-HSA-70268) (Figure 1B, Table 1). For each of the genes in the models, we supported our assertions with standard GO annotation evidence (Balakrishnan *et al*. 2013). 43 assertions were supported by direct experimental evidence from mouse systems, while 16 were supported by sequence orthology to human or rat proteins whose functions are known from experimental data. Genes associated with glycolysis and gluconeogenesis were classified as involved only in gluconeogenesis, only in glycolysis, or shared by both pathways (Figure 2, Table 1).

**Figure 2.**
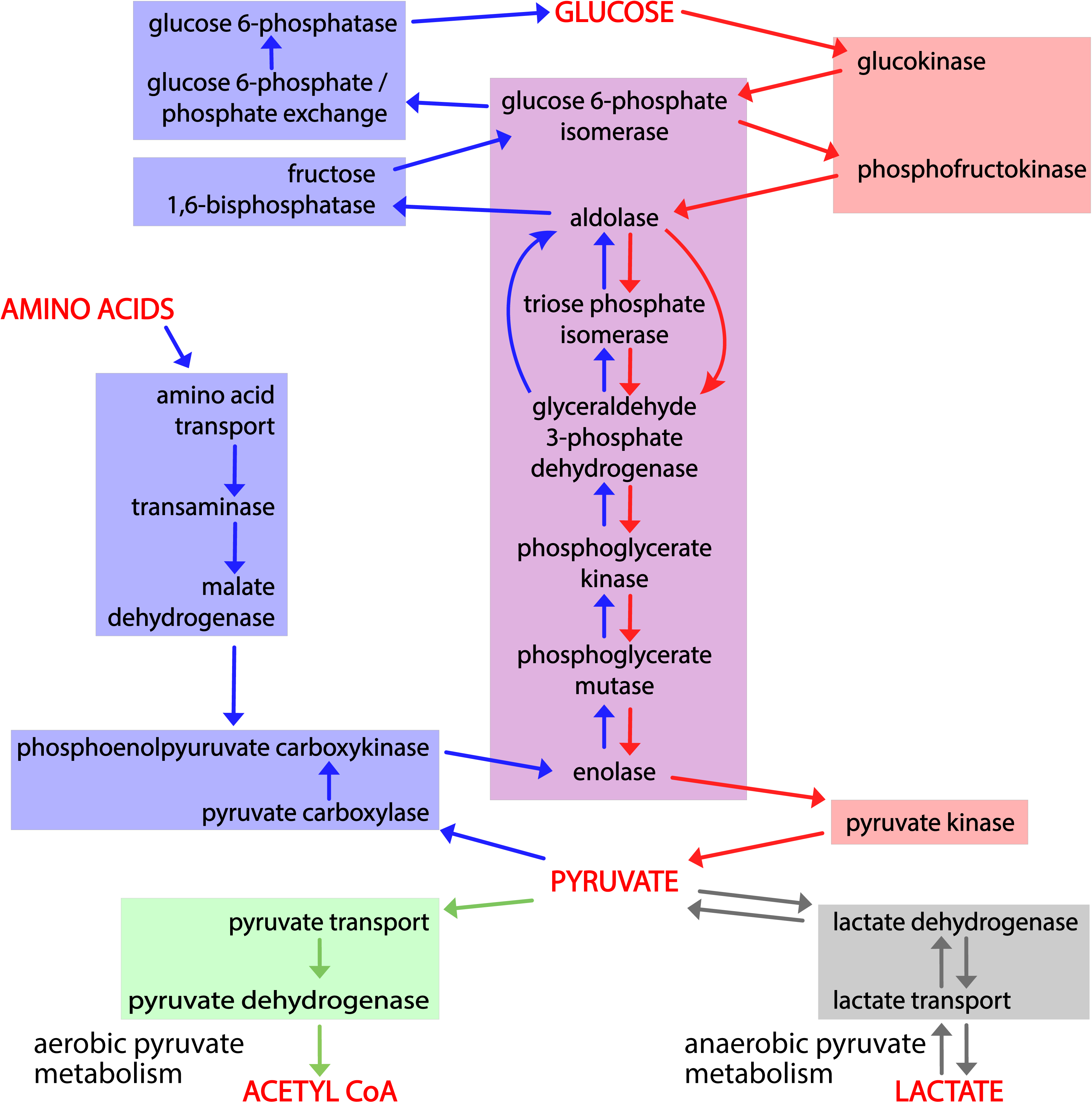
Causal connections among gene products that mediate glycolysis, gluconeogenesis, and pyruvate metabolism. Activity flow representation of the pathways of glycolysis, gluconeogenesis, and pyruvate catabolism, showing gene products (Table 1) associated with each process. The enzymes or transporters that mediate the generation of each required small molecule in these processes are connected by arrows to the ones that consume it. Red arrows connect the steps of glycolysis; blue arrows the steps of gluconeogenesis, and green and gray arrows the steps of aerobic and anaerobic pyruvate metabolism. Red shading highlights enzymes and transporters unique to glycolysis; blue shading enzymes and transporters unique to gluconeogenesis; and purple shading enzymes shared by the two. The green and gray boxes, respectively, indicate the reactions that convert pyruvate to acetyl-CoA or lactate.

### Related Biological Pathways Are Associated with Related but Distinct Sets of Specific Phenotypes

To ask whether variant forms of genes associated with a pathway were associated with similar phenotypes, we used the Mammalian Phenotype (MP) ontology terms assigned by MGD to the genes in our mouse GO-CAM pathways (Table 1) to perform comparative enrichment analyses using the VisuaL Annotation Display (VLAD) tool (Richardson and Bult 2015). 53 of 59 genes in our pathways were annotated with MP terms in MGD. We analyzed three sets of genes, glycolysis only, gluconeogenesis only, and shared between glycolysis and gluconeogenesis. This VLAD enrichment comparative analysis identified 95 phenotypes that were enriched in at least one of the gene sets with an FDR (q) <0.05 (Table s1). The top scoring phenotypes for comparative enrichment, which fall in the domains of homeostasis, embryonic lethality, and hematopoietic system, and their relationships to one another in the MP ontology are shown graphically in Figure 3. The three colors in the bar in each node represent the relative significance values for each gene set. A tabular summary of the relative enrichment data is shown in Table 2, split into three sections: gluconeogenesis only, glycolysis only and shared. The table shows enrichment results for all MP domains that fit three criteria: i) the term has an FDR<0.05, ii) the term has no children that are also FDR<0.05, and iii) when a term or a child passes criteria i) and ii) in more than one set, the terms with the smallest FDR are retained. For example, abnormal hematocrit (shared) is not shown in the table because decreased hematocrit (glycolysis) scores better with these criteria.

**Figure 3.**
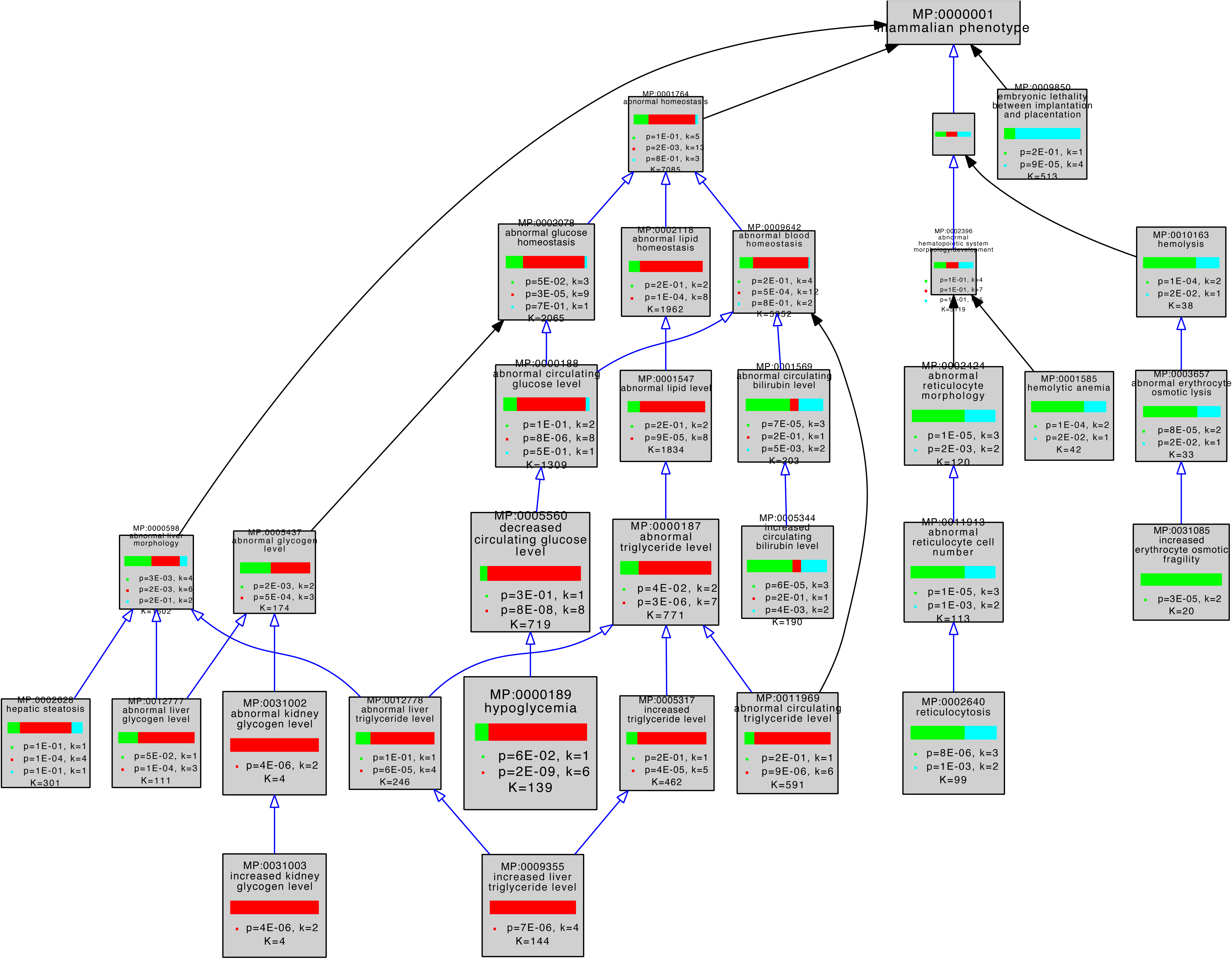
Graphical view of VLAD comparative enrichment analysis for Mammalian Phenotype annotation data from January 26, 2023. Three gene sets were used for the analysis: glycolysis (green), gluconeogenesis (red) and genes shared between glycolysis and gluconeogenesis (blue) (Table 1). Blue arrows indicate a direct is_a relationship between terms, black arrows indicate that the terms are indirectly related. As indicated in Table 1, one glycolysis-only gene (*Pfkp* MGI:1891833) and five shared genes (*Pgk1* MGI:97555, *Pgam2* MGI:1933118, *Eno1b* MGI:3648653, *Eno2* MGI:95394, and *Eno3* MGI:95395) have not been associated with any variant phenotypes and so were excluded from this analysis. Phenotypes in the domains of homeostasis, embryonic lethality, and hematopoietic system are shown. This graph shows the top 25 phenotypes based on enrichment score for clarity. The complete results of this VLAD analysis are tabulated in Table s1.

The three gene sets were associated with distinct enriched phenotypes. The gluconeogenesis genes had more significant enrichment in categories representing ‘abnormal triglyceride levels’, ‘abnormal glycogen levels’, ‘hypoglycemia’, ‘ketosis’ and growth and development. The glycolysis genes had more significant enrichment in categories representing glucose homeostatic phenotypes as well as erythrocyte (in)stability and downstream consequences such as iron levels and bilirubin levels. Shared genes were more significantly enriched for sperm and lethal phenotypes.

The higher levels of the MP ontology hierarchy in Figure 3 show that top scoring abnormal phenotypes for genes unique to glycolysis tend to affect erythropoiesis, while those of gluconeogenesis tend to affect energy metabolism, notably of glycogen and triglycerides.

By convention glycolysis ends with production of pyruvate. Pyruvate is further metabolized aerobically to form acetyl-CoA or anaerobically to form lactate (Figures 1B and 2A) (Prochownik and Wang 2021). Given the distinct phenotypes associated with variation in genes associated with glycolysis and gluconeogenesis we next tested if we could discriminate between phenotypes associated with these two fates of pyruvate. We compared three gene sets: genes that were unique to glycolysis (i.e., not in gluconeogenesis), and these glycolysis genes in combination with those either for aerobic or anaerobic pyruvate metabolism. We analyzed the three gene sets using VLAD and compared phenotypes that were enriched with a FDR <0.05 (Table 3) by counting the number of genes associated with each phenotype.

Phenotypes associated with variant forms of genes of anaerobic metabolism represented abnormalities in the erythropoietic system such as ‘reticulocytosis’ (MP:0002640), ‘hemolytic anemia’ (MP:0001585), ‘decreased hematocrit’ (MP:0000208) and ‘decreased erythrocyte cell number’ (MP:0002875). Phenotypes indirectly associated with the erythropoietic system included ‘increased circulating bilirubin level’ (MP:005344) and ‘abnormal lactate dehydrogenase level’ (MP:002943), two clinical features of hemolytic anemia (Dhaliwal *et al*. 2004). We also observed an increase in the number of genes associated with ‘impaired exercise endurance’ (MP:0012106). One phenotype, ‘decreased circulating alanine transaminase level’ (MP:0002942), was not associated with any genes in the glycolysis pathway, but was associated with two genes, *Ldha* (MGI:96759) and *Slc16a1* (MGI:106013) in the downstream conversion of pyruvate to lactate.

Phenotypes associated with variant forms of genes of aerobic metabolism represented abnormalities such as ‘impaired glucose tolerance’ (MP:0005293), ‘decreased insulin secretion’ (MP:0003059), ‘increased circulating lactate level’ (MP:0013405), and ‘ketosis’ (MP:0030970) (a phenotype associated with rapid lipid breakdown). Several phenotypes with central nervous system defects, e.g. ‘abnormal putamen morphology’ (MP:0004079) and ‘decreased stria medullaris size’ (MP:0020544), were associated with genes in the pyruvate to acetyl-CoA portion of the pathway and not the glycolytic portion. Another phenotype, ‘increased circulating lactate level’ (MP:0013405), was only associated with genes in the downstream pyruvate to acetyl-CoA pathway and may indicate a metabolic switch to the pyruvate pathway when the aerobic pathway is not functional.

We also saw general phenotypes associated with genes that function in both the lactate and acetyl-CoA pathways when compared with core glycolysis, e.g., ‘abnormal triglyceride level’ (MP:0000187), abnormal glucose tolerance (MP:0005291) and ‘decreased circulating insulin level’ (MP:0002727).

### Phenotypes Associated With Specific Pathways Can Be Used to Identify Transcription Factors Causally Related to the Pathway

Since the GO-CAM models we created represent genes that are causally connected in a pathway, we hypothesized that we could make predictions about other genes that might be causally connected to the same pathways based on their phenotypes. Specifically, we attempted to identify DNA binding transcription factors causally connected to gluconeogenesis by searching for transcription factors that shared enriched phenotypes. We used the MGD ‘Genes and Markers’ query tool to search for genes annotated with the Gene Ontology term ‘DNA binding transcription factor activity’ (GO:0003700) and the most specific enriched gluconeogenesis phenotypes with FDR<0.05. The search yielded 93 genes annotated as transcription factors (Table s5), 11 of which were associated with three or more phenotypes (Table 4). We then searched for published evidence of causal connections to gluconeogenesis for each of these 11 genes:

– *Cebpa* (MGI:99480) 6 phenotypes: *Cebpa* directly regulates gluconeogenesis by regulating the gluconeogenic enzymes phosphoenolpyruvate carboxykinase and glucose-6-phosphatase (Pedersen *et al*. 2007). Furthermore, knocking out *Cebpa* (*Cebpa^tm1Gjd^*, MGI:2177053) results in ‘abnormal gluconeogenesis’ (MP:0003383) (Wang *et al*. 1995).
– *Bmal1* (MGI:1096381) 5 phenotypes: *Bmal1* regulates gluconeogenic gene expression by interaction with *Hdac5* (Li *et al*. 2018). *Bmal1* interacts with *Clock* to form a transcription factor complex (Huang *et al*. 2012) involved in the circadian regulation of glucose homeostasis (Schiaffino *et al*. 2016).
– *Cebpb* (MGI:88373) 4 phenotypes: *Cebpb* directly regulates the expression of the phosphoenolpyruvate carboxykinase gene (Arizmendi *et al*. 1999).
– *Pparg* (MGI:97747) 4 phenotypes: The *Pparg* gene is generally thought to have a negative effect on gluconeogenesis and is involved in adipocyte differentiation (Pendse *et al*. 2010); (Hernandez-Quiles *et al*. 2021); (Way *et al*. 2001). However *Pparg* may indirectly upregulate gluconeogenesis through *Pomc* signaling in response to a high fat diet (Long *et al*. 2014). The *Pparg2* isoform also upregulates pyruvate carboxylase gene expression (Jitrapakdee *et al*. 2005). Three of the four phenotypes associated with *Pparg* involve triglyceride metabolism. These could all reflect its overall involvement in controlling triglyceride metabolism (Semple *et al*. 2006), consistent with the fourth ‘hepatic steatosis’ phenotype.
– *Ncoa3* (MGI:1276535) 3 phenotypes: We were not able to find direct evidence of a role for this gene in gluconeogenesis. It is well-documented in the control of lipid metabolism (Astapova 2016) and members of the *Ncoa* family are thought to be overall regulators of metabolic processes (York and O’Malley 2010).
– *Foxa2* (MGI:1347476) and *Foxo1* (MGI:1890077) 3 phenotypes each: *Foxa2* in concert with *Foxo1*, upregulates gluconeogenesis and in particular upregulates phosphoenolpyruvate carboxykinase in the fasting liver (Puigserver and Rodgers 2006; Zhang *et al*. 2005).
– *Klf15* (MGI:1929988) 3 phenotypes: *Klf15* is one of the genes responsible for regulating the switch downregulating lipogenesis and upregulating gluconeogenesis during fasting, upregulating the expression of *Pck1*) (Takeuchi *et al*. 2016). Additional results support the role of *Klf15* in regulating gluconeogenesis through mutational studies and transcription assays (Gray *et al*. 2007; Teshigawara *et al*. 2005).
– *Nr1h4* (MGI:1352464) 3 phenotypes: Metabolites of sesame oil antagonize Nr1h4 function, resulting in a decrease in mRNA levels encoding phosphoenolpyruvate carboxykinase and glucose 6-phosphatase (Sasaki *et al*. 2022). Activation of Nr1h4 by cholic acids, however, resulted in a decrease in the expression of gluconeogenic genes, likely by indirectly regulating Foxo1 and Hnf4a (Yamagata *et al*. 2004). These results may be a reflection of the interaction of Nr1h4 with different partners, resulting in different outcomes (Yamagata *et al*. 2004; Xu *et al*. 2018).
– *Ppara* (MGI:104740) 3 phenotypes: *Ppara* plays a role in regulating many metabolic genes including those that function in gluconeogenesis and lipid metabolism (Preidis *et al*. 2017). Like other transcription factors we have identified in our query, the exact nature of *Ppara*’s activity may depend on its partners and other transcription factors that are expressed in different physiological states (Kersten 2014).
– *Clock* (MGI:99698) 3 phenotypes: *Clock* and *Bmal1* (above) interact to synchronize gluconeogenesis with circadian rhythms(Li *et al*. 2018; Schiaffino *et al*. 2016).

Although it is standard practice at MGD to infer the biological processes in which a gene is involved using the ‘inferred from mutant phenotype’ evidence code, GO and phenotype annotation are done separately. Of the 11 genes that we identified through our phenotypic analysis, only two, *Ppara* and *Foxo1*, were already annotated in the separate GO curation process as being involved in gluconeogenesis or its regulation. Six of the 11 transcription factors we identified (*Cebpa*, *Bmal1*, *Cebpb*, *Foxo1*, *Klf15*, *Clock*) were also annotated to ‘abnormal gluconeogenesis’ (MP:0003383). To test whether the transcription factors were preferentially annotated to gluconeogenic phenotypes, we ran a VLAD analysis on the transcription factors alone and compared with the gluconeogenic genes. As expected, the top scoring phenotypes were related to the pre-chosen gluconeogenic phenotypes. However, the transcription factors were also significantly enriched for other phenotypes that were not enriched in the gluconeogenic gene analysis, eg. ‘abnormal ovulation cycle’ (MP:0009344) and ‘abnormal hair cycle anagen phase’ (MP:0008858) (supplemental table S6, analysis date July 3, 2023). These results imply that the transcription factors we identified were not *a priori* or specifically limited to the study of gluconeogenesis.

## Discussion

Here, we have shown that human GO-CAM models of glycolysis, gluconeogenesis, and pyruvate metabolism serve as accurate templates for modeling the orthologous mouse pathways, that VLAD analysis of phenotypes associated with mutant mouse genes associated distinct but related phenotypes with mutational disruptions of genes involved in each pathway, and that analysis of common enriched phenotypes further supported the identification of transcription factors that play major roles in regulating these pathways. These results are not surprising: these pathways, their transcriptional regulation, and their pathophysiology have been very extensively characterized experimentally in both species and shown to be closely related. Neither GO-CAM construction nor VLAD analysis was trained or optimized for these domains of biology, however, so these results are useful as a test of a general strategy for pathway annotation and analysis, and suggest that it can be extended to less exhaustively studied processes and less well characterized species.

### Reactome GO-CAMs can bridge the biology of model organisms and humans

Reactome annotations of human pathways of glycolysis, gluconeogenesis, and pyruvate metabolism to lactate or acetyl-CoA identified functions for 60 human gene products, 44 of which have high-confidence mouse structural orthologs (Table1), enabling the streamlined generation of mouse GO-CAM models by swapping human gene products for their mouse counterparts. This successful manual pathway construction exercise is a proof of principle for automating the construction of model organism template pathways useful as starting points for expert manual curation and addition of organism-specific supporting experimental evidence. Indeed, Reactome already computationally predicts pathways for fifteen model organisms using orthologs of human proteins to systematically populate reactions for these other species, a process that could be extended to any model organism whose proteome is known (https://reactome.org/documentation/inferred-events).

### Phenotype enrichment analysis of pathway-specific gene lists identifies and distinguishes pathway-specific phenotypes

We separated the genes annotated in our mouse GO-CAM models into three groups: specific for glycolysis, specific for gluconeogenesis, and shared genes. Phenotype enrichment on the three sets showed that the enriched phenotypes fell into different classes for each set (Figures 2 and 3). Phenotypes associated with alleles of the genes shared by glycolysis and gluconeogenesis compared with genes specific to each pathway were overrepresented in an early embryonic lethal phenotype, reflecting the central role of glucose metabolism in mammalian physiology.

Variants of the glycolysis-specific genes, compared to gluconeogenesis-specific or shared genes, were more significantly enriched for phenotypes associated with abnormalities of red blood cells, consistent with the absolute dependence of these cells on anaerobic glycolysis for ATP.

Variants of gluconeogenesis-specific genes were overrepresented in the abnormal phenotypes of hypoglycemia, increased triglyceride levels and abnormal kidney and liver glycogen levels. Increased glycogen levels in the kidney and liver are explained by the identities of the genetic defects associated with the phenotypes: mutations that inactivate *Slc37a4* (MGI:1316650) and *G6pc* (MGI:95607). The products of these two genes normally mediate the transport and dephosphorylation of glucose-6-phosphate newly synthesized by gluconeogenesis, enabling release of free glucose into the circulation – inactivation of either leads directly to hypoglycemia (Froissart *et al*. 2011). Abnormal accumulation of intracellular glucose-6-phosphate drives glycogen synthesis by mass action (Adeva-Andany *et al*. 2016). ‘Increased liver triglyceride level’ was associated with mutations of three genes in the gluconeogenesis-specific gene set, *G6pc* (MGI:95607), *Pck1* (MGI:97501), and *Slc25a13* (MGI:1354721), and is consistent with phenotypes associated with excess accumulation of glucose-6-phosphate observed in a *G6pc* knockout mouse and in human patients with partial loss of G6PC function (Hoogerland *et al*. 2021).

To test the hypothesis that phenotypes could be further distinguished by joining analysis of glycolysis with analysis of the aerobic and anaerobic fates of its major product, pyruvate, we performed phenotype enrichment analyses on three sets of genes: those specific for canonical glycolysis (conversion of glucose to pyruvate), those for canonical glycolysis plus downstream anaerobic metabolism of pyruvate to lactate, and those specific for canonical glycolysis plus downstream aerobic metabolism of pyruvate to acetyl-CoA (Table 3). Analysis of the glycolysis + lactate/anaerobic gene set revealed additional associations of genes with red blood cell phenotypes. These phenotypes reflect the dependence of red blood cells on anaerobic glycolysis to lactate as an energy source (Goto *et al*. 2019); (Rose and Warms 1966); (TeSlaa *et al*. 2021). A non-erythropoietic phenotype identified in this analysis was ‘impaired exercise endurance’, consistent with enhanced anaerobic glucose metabolism during intense exercise (Alberti 1977; Robergs *et al*. 2004). When we add the genes for the acetyl-CoA/aerobic pathway, additional associations with glucose homeostasis phenotypes are detected. These associations are in concordance with known physiology. Glycolysis and the downstream aerobic metabolism of glucose through pyruvate conversion to acetyl-CoA and the tricarboxylic acid cycle are key steps in the regulation of insulin secretion in response to glucose in pancreatic beta cells (Newgard and McGarry 1995).These results show that we can identify connected pathways that give rise to distinct phenotypes and to distinguish subnetworks in a branching system that have discreet biological outcomes.

### Enriched Phenotypes can be used to identify genes that are causally connected to a pathway

To test our ability to identify genes that are not conventionally associated with a pathway but that are causally connected to it by shared phenotypes, we used enriched phenotypes in the gluconeogenesis pathway to search for transcription factor genes whose variants were associated with these same phenotypes. *Bmal1* (Li *et al*. 2018) *Cebpa* (Pedersen *et al*. 2007), *Cebpb* (Arizmendi *et al*. 1999) and *Foxa2* (Puigserver and Rodgers 2006; Zhang *et al*. 2005), either have a direct or indirect effect on the expression of gluconeogenesis genes. Several genes, such as *Ncoa3* (York and O’Malley 2010) and *Pparg* (Semple *et al*. 2006) are also connected in less direct ways to gluconeogenesis and may be connected to other pathways causally connected to gluconeogenesis such as glucagon signaling and lipid metabolism. Some genes, like *Ppara*, regulate gluconeogenesis as well as other related, connected metabolic processes like lipid metabolism (Rakhshandehroo *et al*. 2007). Our results are consistent with a complex regulation of gluconeogenesis where a variety of transcription factors act together both directly and indirectly to coordinate metabolism depending on specific contexts (DeSousa-Coelho *et al*. 2023; Li *et al*. 2018; Pei *et al*. 2006; Qu *et al*. 2021).

In an extension of previous work demonstrating that phenotypes can be used to predict GO annotations, our work further shows the integrated nature of phenotypic analysis and functional inference (<u>Ascensao</u> *et al*. 2014). With the exception of two transcription factors, *Ppara* and *Foxo1*, none of the genes we identified by shared phenotypes were yet annotated to gluconeogenesis GO terms even though six of them were annotated to the phenotype term ‘abnormal gluconeogenesis’ (MP:0003383). Although it is serendipitous that some of the transcription factors had not yet been annotated to GO based on phenotypes associated with gluconeogenesis, this suggests that genes were not ‘selected’ with a pre-existing annotation focus on gluconeogenesis. Instead, it shows that phenotypes can be used to inform us about gene function and that gene function may be useful for inferring phenotypes.

We hypothesize that as we integrate models of additional regulatory processes with models of metabolism, we will be able to discriminate among the networks that these transcription factors modulate and identify specific gene targets within them. Concatenation of regulatory and housekeeping processes will enable construction of vast network hairballs; the limited study described here suggests that it may be possible to manage and interpret them to generate integrated models of cell physiology useful for modeling and experimental design.

### Limitations of our current approach and future directions

To increase the sensitivity of our search for abnormal phenotypes associated with pathways, we have grouped the effects of all paralogs that normally mediate each activity in glycolysis, gluconeogenesis, and pyruvate metabolism. Despite this grouping, known tissue– and process-specific effects, such as the impact of glycolysis defects on red cell function and sperm cell maturation and function, and of gluconeogenesis defects on glucose homeostasis can be detected above background significance levels (Table s1). Nevertheless, we have missed other known effects, such as the opposing effects of variant forms of GCK (human) / Gck (mouse – MGI:1270854) that have abnormally high and low affinities for glucose on insulin secretion in response to varying blood glucose levels. This result suggests that our analysis could be refined further by taking into account the nature of the alleles that result in mutant phenotypes, the background strains on which alleles are present, and the physiological conditions in which phenotypes are measured. We also failed to observe red blood cell phenotypes for all genes required for conversion of glucose to pyruvate. These failures have several possible explanations. First, the collection of known genetic variants is far from saturating, and phenotypic characterization of existing variants has not been exhaustive. Second, many variants in these genes are likely to be incompatible with early embryonic development, so their effects on postnatal functions would not be detectable. Third, most of the steps of glycolysis are mediated by sets of paralogous genes that differ from one another in their regulation and their tissue-specific expression, but it is plausible that phenotypes due to variants in one member of a set could be masked by even low level expression of other (wild-type) set members.

As in all bioinformatics approaches, our analyses are limited by the data available. We chose well-studied pathways for this work to test whether when the data exist, we can use those data to generate specific conclusions. In cases where phenotypic data are not available or comprehensive for a large number of genes in a pathway or the pathway is not well-defined, our approach will be limited due to lack of enrichment or multiple genes showing the same phenotype and phenotypes may be missed. There will be a ‘tipping point’ in which enough genes in a pathway will be annotated to a phenotype to result in a significant enrichment or will give a clear indication that disturbing the pathway results in a phenotype. However, once the ‘tipping point’ is reached, we can predict that perturbing the other genes in the causal chain will result in the same phenotype.

Relating biochemical pathways to phenotypes associated with specific genes provides a tool to predict and test the effects of genetic or pharmacological manipulation on the physiology of an organism. Identifying the elements of pathways, including ones with complicated structures, that result in a phenotype should better support targeted approaches to modify the pathways and change phenotypic outcomes. Beyond the classic approach of providing exogenous sources of key molecules missing due to a gene defect, we can envision strategies to manipulate pathways that are hypermorphic and are deleterious to an organism, with pharmacologic inhibitors or RNAi methods. Finally, these types of analyses may help us to dissect the complex interactions of drugs with biochemical pathways to better understand mechanisms and potential side effects that could be addressed with co-therapies. For example, metformin is one of the primary pharmaceuticals used to treat type-2 diabetes. Metformin lowers gluconeogenesis in the liver, but its actions are complex and not completely understood (Rena *et al*. 2017; Flory and Lipska 2019). As we further enrich our representation of glucose metabolism to include metabolic and regulatory networks, it might be possible to perform a detailed analysis of the effects (phenotypes) of metformin treatment in mice and compare that to network analysis similar to the ones we performed in this work. The comparison can be used to identify potential branches in the network where metformin acts by looking for shared or unique phenotypes.

The available mouse phenotyping data themselves present some limitations. Different mutations in the same gene can have variable phenotypes due to the molecular nature of the mutation (e.g., null or knockout vs. point mutation, or isoform-specific mutations). Mutations may be hypomorphic, gain or loss of function or even silent in the absence of an inducer or stressor. Phenotypic variability due to strain-specific variants in the mouse experimental genetic background can contribute to conflicting results when the data is inferred at the gene level (Perry *et al*. 2020). Genetic sex, microbiota and parental origin (imprinting) can also influence phenotypes. Outliers in the data should be assessed for these potential causes in variability.

In summary, we have shown that we can convert a set of human biochemical pathways to corresponding pathways in the laboratory mouse. We can then take advantage of the rich genetic work that has been done in the mouse to analyze and determine the phenotypic consequences of perturbations of genes in those pathways. In this way, we integrate the information about human biology from Reactome with model-organism biology from MGD. We envision that this approach can be extended to other metabolic processes and other model organisms and the study of pathway-phenotype connections can be used not only to understand the similarities of the pathways but as a testing ground for manipulation of pathways in more experimentally tractable organisms than human.

## Acknowledgements

We dedicate this paper to the memory of Michael Ashburner. The work described here comes out of projects he helped to initiate and guide and we benefitted immeasurably from his support.

This work was supported by NIH grants U41 HG002273 and U24 HG012212 (GO Consortium), U24 HG012198 (Reactome), U24 HG002223 (WormBase), U41 HG000330 (The Mouse Genome Database), and U24 HG011851 (Pathways2GO).

---

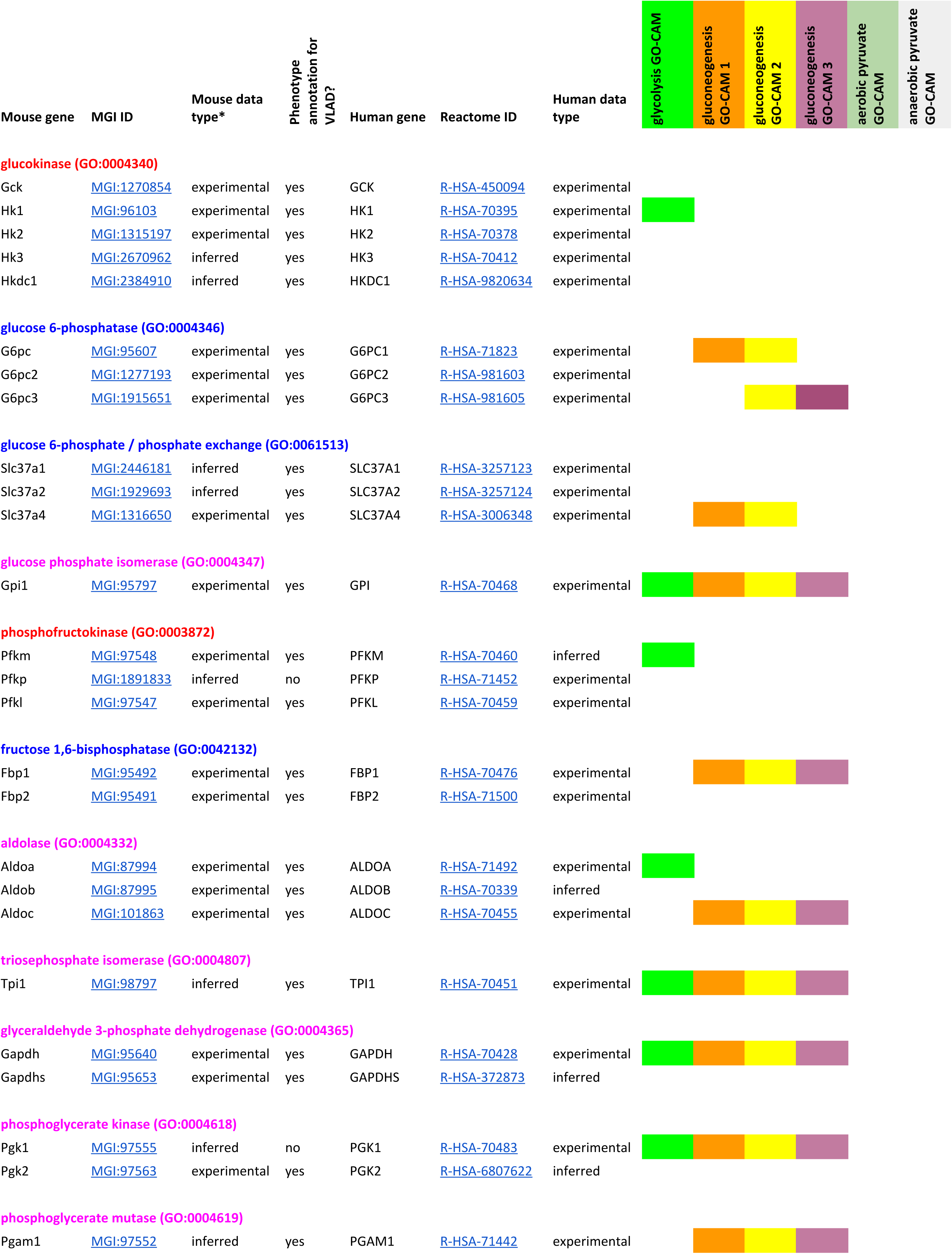
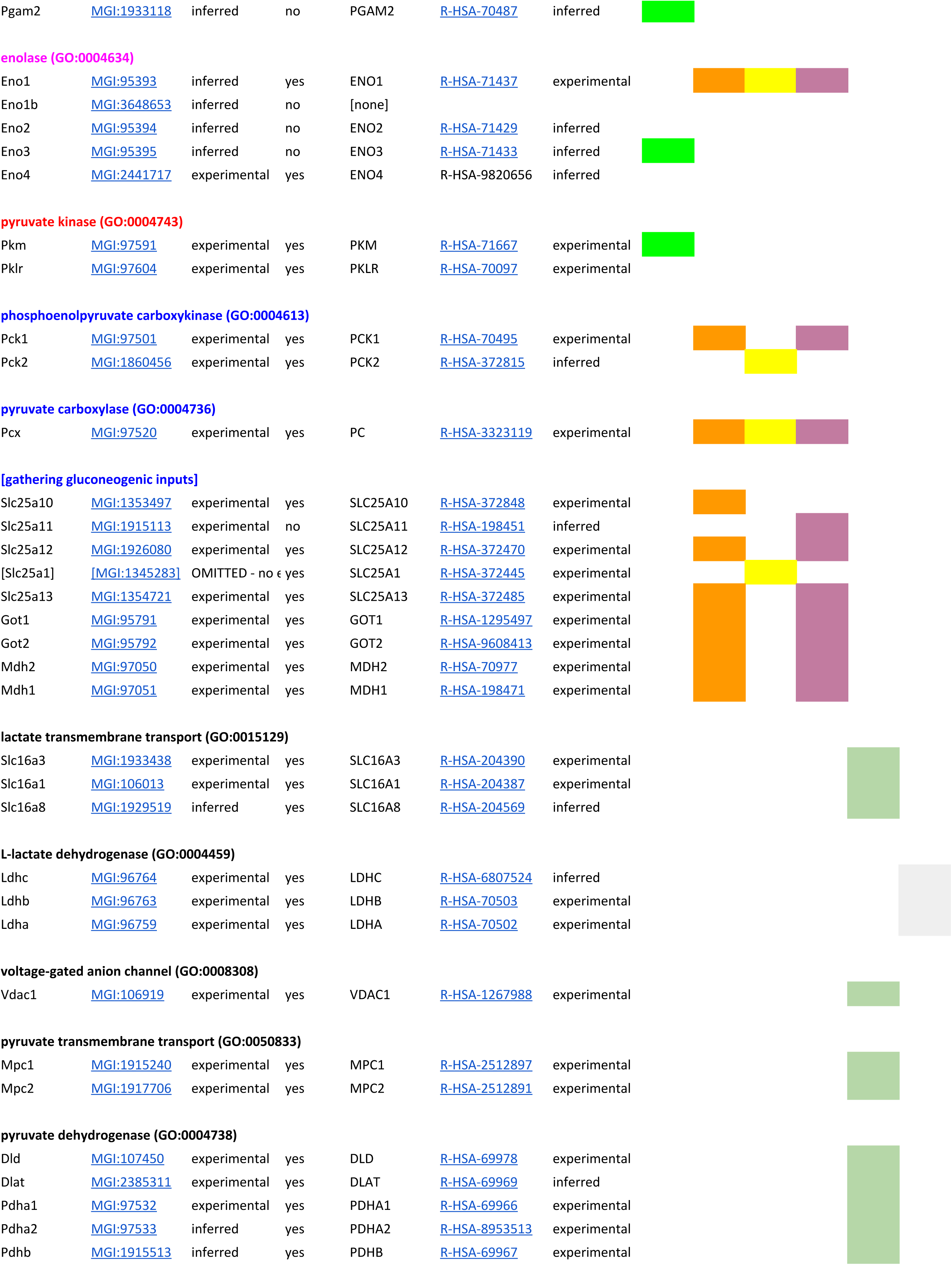
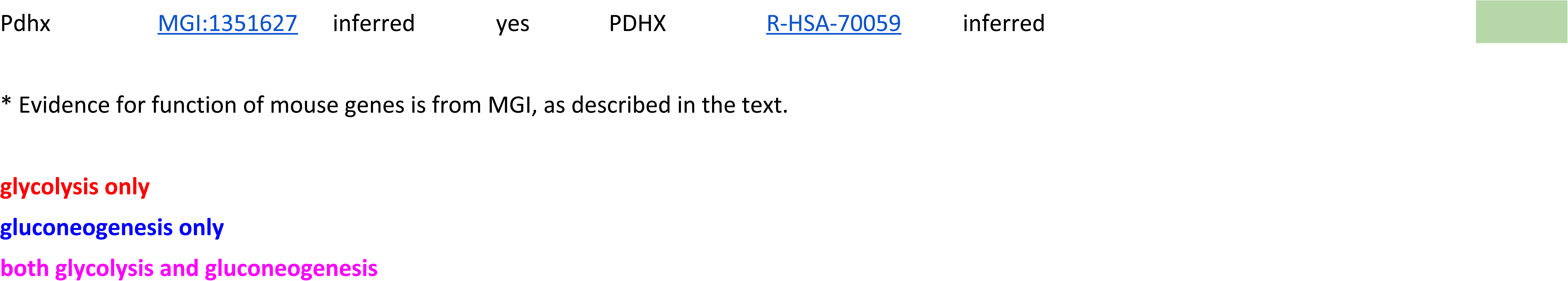
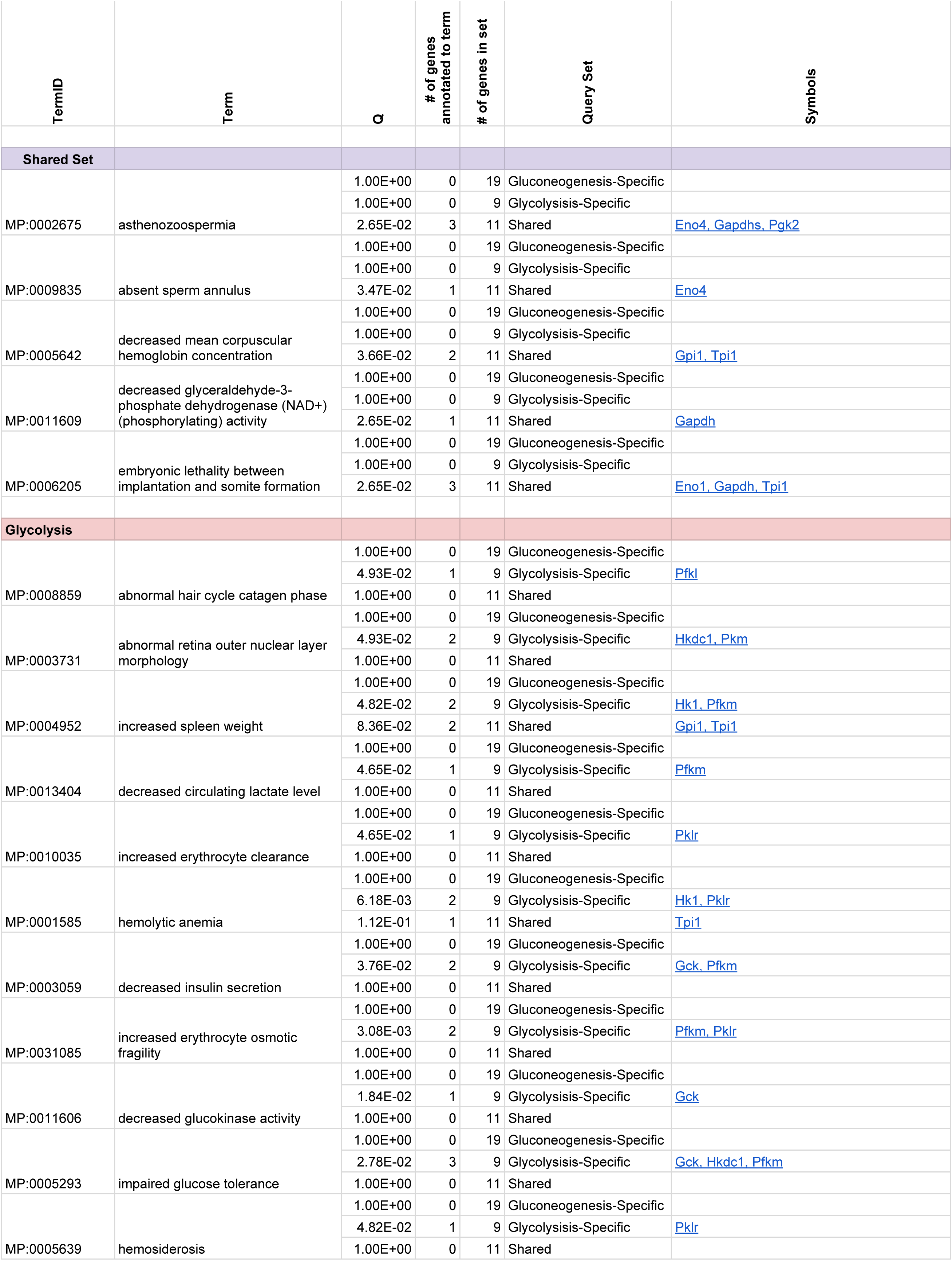
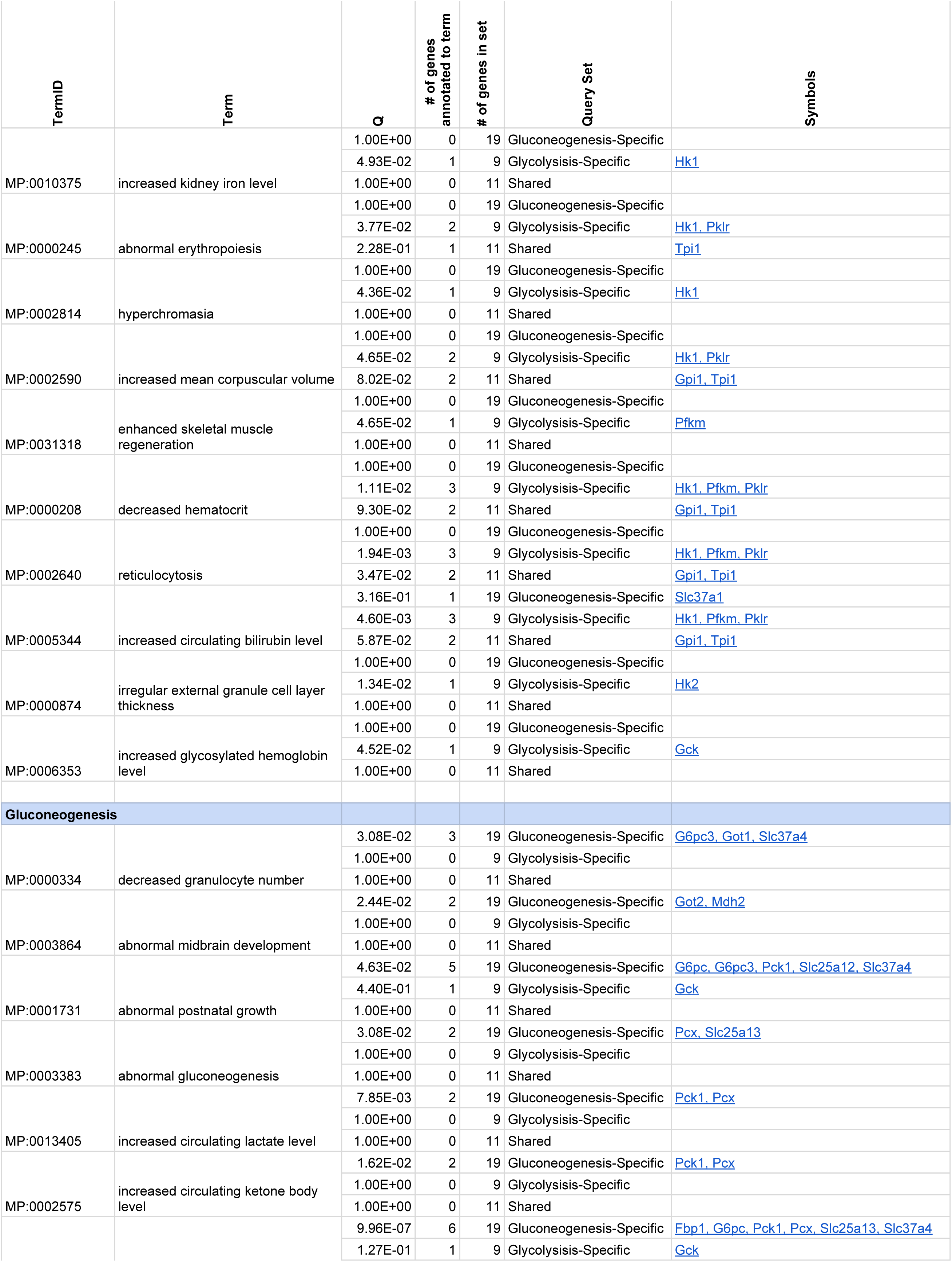
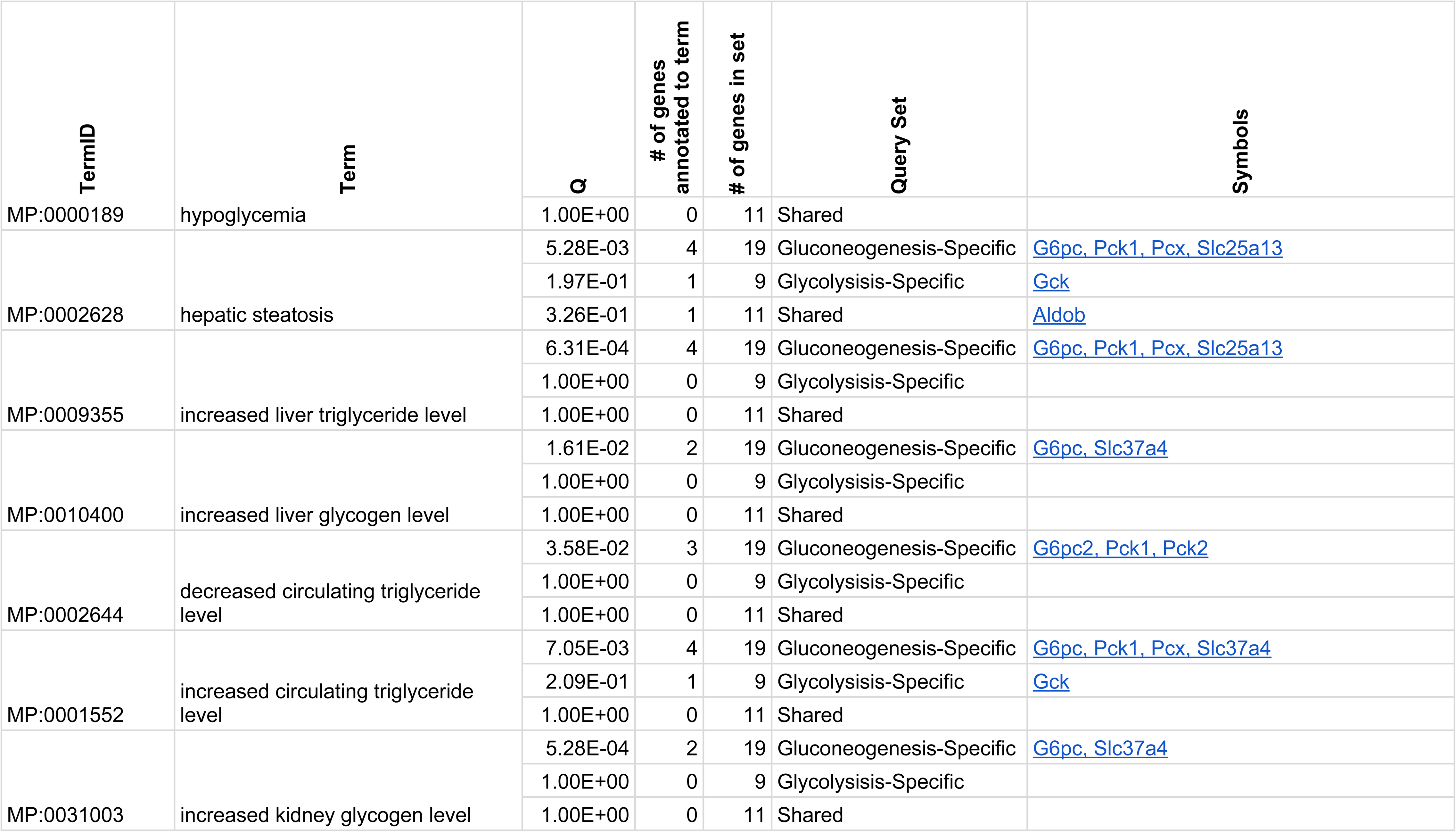
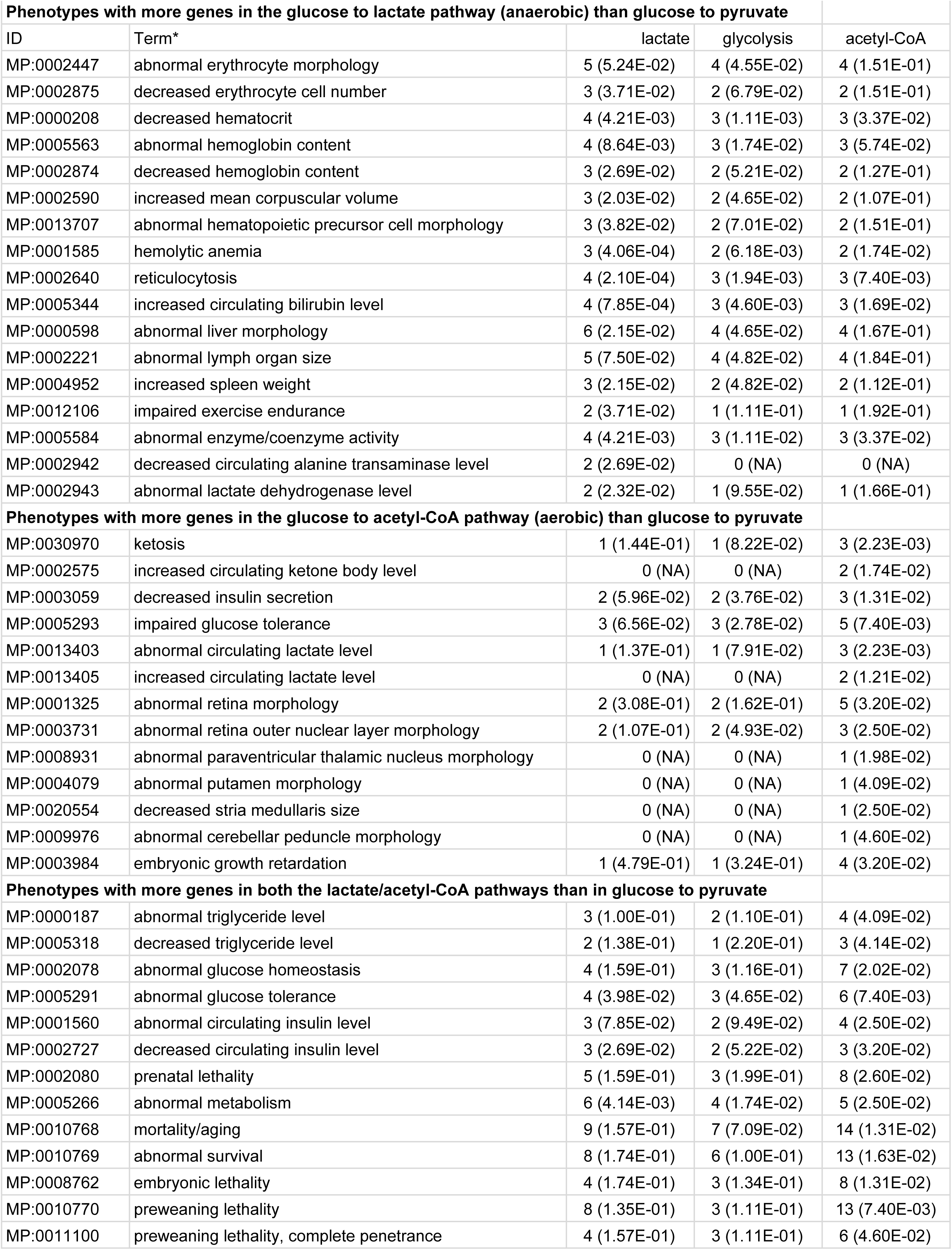
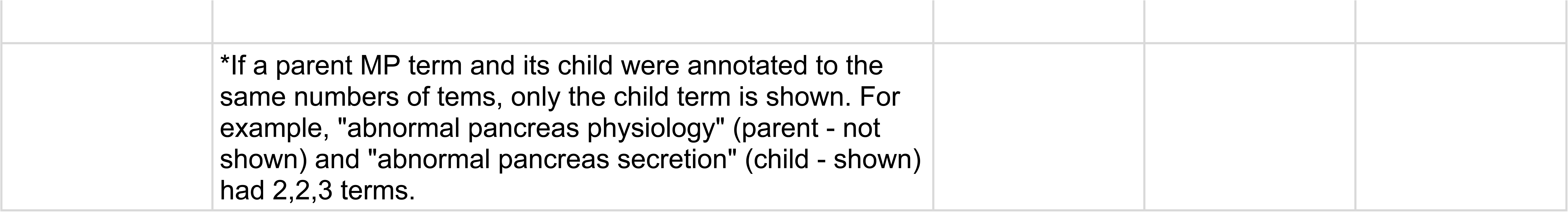
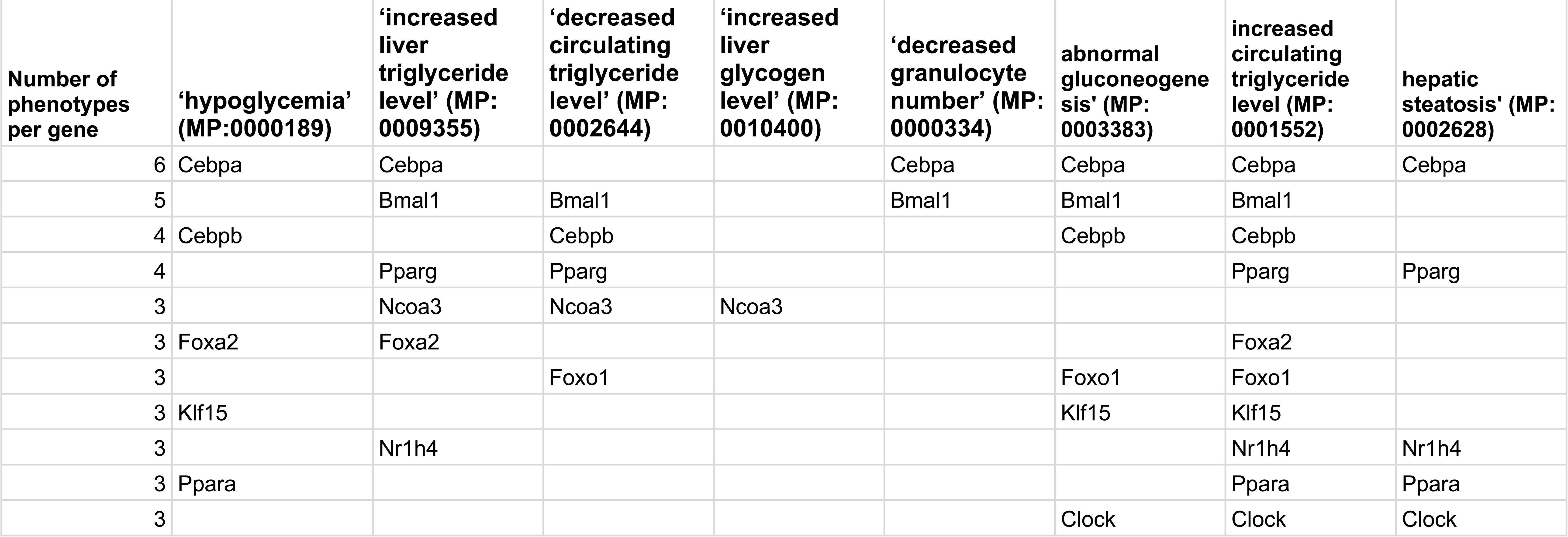

